# Evaluation of plasma anti-CS3 and anti-LTB IgG avidity among Zambian children vaccinated with ETVAX

**DOI:** 10.1101/2025.10.12.681928

**Authors:** Cynthia Mubanga, Mutale Mubanga, Obvious Nchimunya Chilyabanyama, Masauso Phiri, Caroline C. Chisenga, Richard H. Glashoff, Roma Chilengi

## Abstract

**Background:** Enterotoxigenic *Escherichia coli* (ETEC) remains a major cause of diarrheal disease in low- and middle-income countries (LMICs). To curb ETEC related diarrhoea, several candidate vaccines are in development, with ETVAX^®^ being the most advanced. Although immunogenicity studies have primarily focused on measuring antibody titres, assessing antibody avidity offers additional valuable insight into antibody quality and immune maturity. This study assessed anti-CS3 and anti-LTB IgG avidity in Zambian children to better understand vaccine-induced antibody responses in an endemic setting.

**Methods:** Children aged 6–23 months (n=60) received three quarter-doses of ETVAX^®^ with dmLT adjuvant on days 0, 14, and 90. Plasma samples collected at baseline (V1), seven days after the second dose (V5), and seven days after the third dose (V7) were analysed by limiting antigen dilution ELISA to calculate avidity indices (AI). Naïve classification was performed using titre-based thresholds (20th percentile of baseline titres) and avidity-defined naivety (AI < 0.5). Receiver operating characteristic (ROC) analysis was used to evaluate the discriminatory performance of avidity indices against titre-defined naïve status.

**Results:** Baseline avidity was detectable for both CS3 and LTB, consistent with prior natural exposure. Mean CS3 IgG avidity decreased from 0.7 at baseline to 0.6 after the third dose (*p*<0.001), while LTB IgG avidity showed transient decreases but no net gain. Naïve classification at baseline revealed that 9/60 children had titres but low avidity (functional naivety), and 6/60 had waned titres but high avidity. Only one child was naïve by both criteria for CS3, and none for LTB. ROC analysis demonstrated moderate discrimination for CS3 (AUC=0.65; optimal cut-off AI=0.36) but poor discrimination for LTB (AUC=0.30).

**Conclusion:** In this endemic population, ETVAX^®^ induced strong antibody titres but minimal changes in avidity over time with inter-individual differences, while ROC analysis highlighted the need for context-specific thresholds. These findings show the need for both antibody titre and avidity assessment in vaccine evaluations in endemic settings.

## Background

Diarrheal disease remains a significant global health challenge accounting for about 1.7 million deaths in 2021, with resource-limited settings bearing the largest burden of the disease due to poor access to clean water and inadequate sanitation [1]. Enterotoxigenic *Escherichia coli* (ETEC) is one of the main causes of diarrhoea among children under five years of age in low- and middle-income countries (LMICs), significantly contributing to morbidity and mortality [1–3]. Globally, ETEC is responsible for an estimated 220 million diarrheal episodes annually, with 75 million occurring in children under five [4]. Although ETEC-induced diarrhoea is often asymptomatic or self-limiting, it can cause gut inflammation, leading to risks such as environmental enteric dysfunction (EED) and malabsorption syndrome, which result in growth impairment and malnutrition that have long-term adverse effects on the children’s development [4, 5].

ETEC pathogenesis begins with bacterial attachment to enterocytes via colonization factors (CFs), followed by the release of heat-stable or heat-labile enterotoxins, leading to watery diarrhoea [6, 7] Protective immunity develops through the generation of antibodies targeting various CFs, which help prevent infection by homologous strains [8]. Over time, affinity maturation enhances antibody binding strength, a process that occurs in the germinal centres of lymphoid tissues with support from T-helper cells and follicular dendritic cells [9, 10]. This maturation is closely linked to antibody class switching (from IgM to IgG, IgA, or IgE) and the differentiation of B cells into plasma cells or memory B cells [9, 11]

Antibody avidity, which reflects the overall binding strength of an antibody to its target antigen, serves as a key indicator of functional humoral immune maturation [12]. As avidity increases with repeated exposure to infection or vaccination, it signals the development of effective immunological memory with high-avidity antibodies forming stable complexes that strongly neutralize antigens, reinforcing long-term immunity [9, 11]. Therefore, antibody avidity is a crucial parameter in immunological studies, providing valuable information for understanding immune response dynamics and the development of immune memory [9].

ETVAX^®^, an advanced candidate vaccine against ETEC, comprises an inactivated *E. coli* bacteria overexpressing colonization factors CFA/I, CS3, CS5, and CS6, along with LCTBA (A hybrid protein between the B-subunit of *E.coli* heat-labile enterotoxin (LTB) and the B-subunit of the cholera toxin (CTB)) and the dmLT (double mutant heat-labile toxin) adjuvant [13]. Clinical trials have shown promising results, demonstrating both safety and immunogenicity, including in children in Bangladesh and Zambia [13, 14]. The phase 1b trial conducted in Zambia found that a quarter dose of ETVAX^®^ combined with the mucosal adjuvant dmLT was immunogenic in children aged 6-23 months, leading to its current evaluation in a phase 2 trial in The Gambia [13].

Plasma anti-CFA/I, CS3, CS5, CS6, and LTB IgA, along with anti-LTB IgG titres, were measured and results showed significant increases in plasma IgA and IgG titres in children who received the ¼ dose after two and three doses compared to placebo[13]. The 6–9-month-old group exhibited a greater magnitude of response than the 10–23-month-olds, likely due to their being more immunologically naïve, as reflected by the higher pre-vaccination titres in the older cohort [13].

To further understand the maturation of the humoral immune responses and the development of immune memory in Zambian children, we investigated changes in antibody avidity following vaccination among participants who received a quarter dose of the ETVAX^®^ vaccine as this dose induced the best titres.

## Methods

### Study design

This was a laboratory-based study that used samples collected during a phase 1 trial aimed at evaluating the safety, tolerability, and immunogenicity of the ETVAX^®^ vaccine with mucosal adjuvant dmLT along with dose selection. The trial participant recruitment took place from 21^st^ September 2019 to 10^th^ September 2020, beginning with enrolment of healthy adults aged 18- 45 years (cohort A). Subsequently, the trial progressed to include children aged 10-23 months (cohort B), followed by those aged 6-9 months (cohort C) once safety was established by the Data Safety and Monitoring Board (DSMB) **(Fig 1).** In cohort A, participants received either one full dose of ETVAX^®^ + 10 µg dmLT or a placebo. Cohorts B and C participants received three doses of either 1/8 or 1/4 of the full ETVAX^®^ dose + 2.5 µg dmLT, or placebo, administered on Days 1, 15, and 90 on an outpatient basis **(Fig 1).**

**Fig 1:**
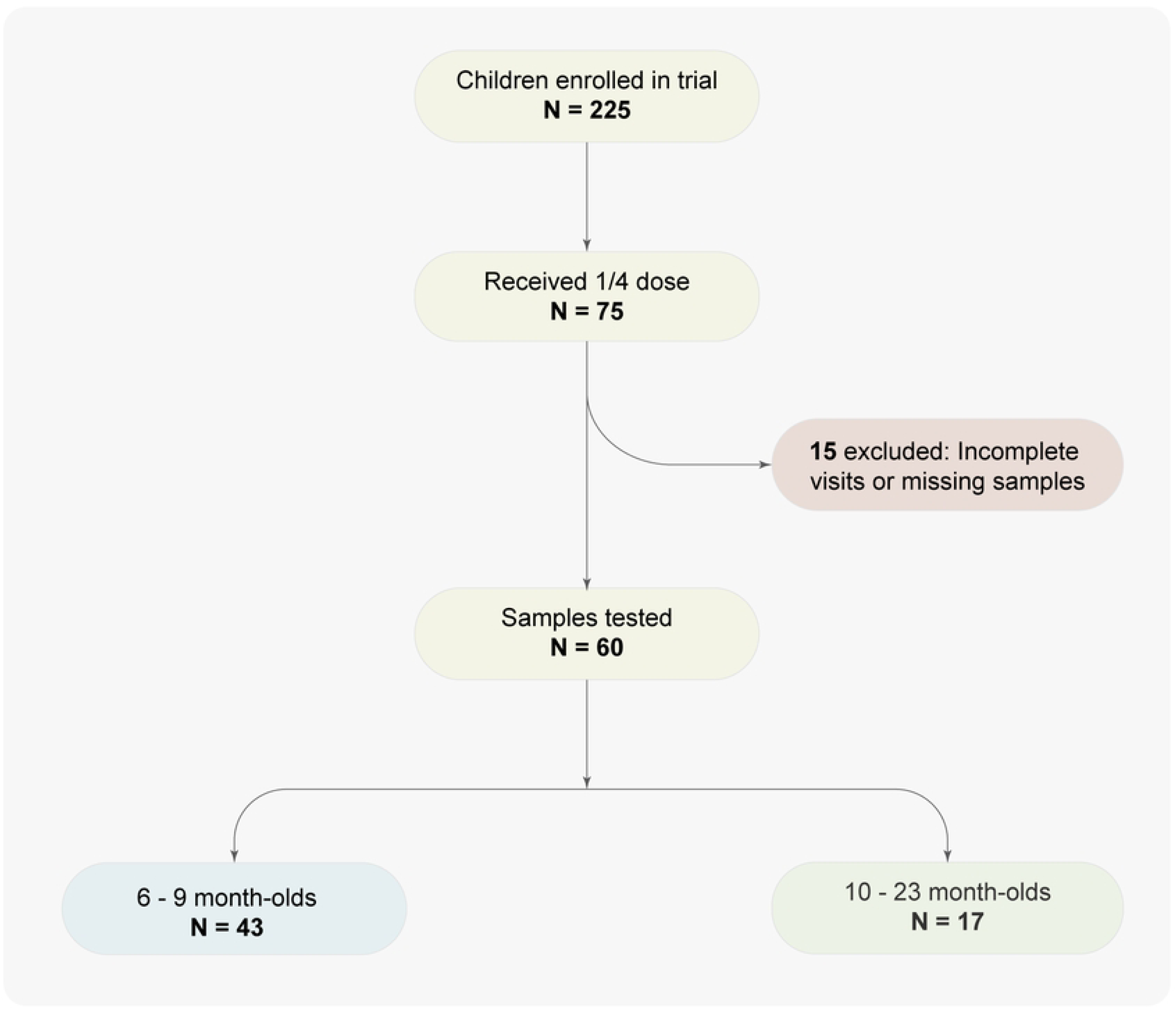
Participant selection flow for avidity analysis. Out of the eligible 75 quarter dose participants, 60 provided complete samples for inclusion.

Ethical approval was obtained from the University of Zambia Biomedical Research Ethics Committee (UNZABREC); reference number 965-2020 to allow for use of stored clinical trial samples. The clinical trial protocol was also approved by UNZABREC, the National Health Research Authority (NHRA), and the Zambia Medicines Regulatory Authority (ZAMRA), and was registered with the Pan African Clinical Trials Registry (PACTR); trial number PACTR201905764389804.

Mothers and caregivers for each participant provided written informed consent before the initiation of any study procedures.

### Specimen collection

Blood samples were collected at baseline, prior to the first vaccination, at seven days after the second dose (Day 22 or V5) and again seven days after the third dose (Day 97 or V7). For this analysis, samples from participants in cohort B and C who received a quarter dose of ETVAX^®^ and attended all study visits, ensuring blood collection at all three time points (Days 1, 22, and 97) were identified. A total of 60 participants (17 from cohort B and 43 from cohort C) out of the 75 that received the quarter dose met these criteria and were included in the study.

### ELISA Testing and Antibody Avidity Assessment

The limiting antigen dilution (LAD) ELISA method, previously described by Leach et al. was employed to measure antibody avidity [6]. The antibody avidity was determined by assessing the specific antibody binding (determined by endpoint titres via serial dilutions) to two different antigen coating concentrations (LTB: 0.1 and 0.02 µg/ml, CS3: 0.3 and 0.06 µg/ml) and calculating the avidity index (AI = titre at lower concentration / titre at higher concentration) [6]. In brief, 96-well ELISA plates (Greiner 655061 for CS3 and NUNC 269620 for LTB) were coated with purified ETEC antigens provided by Gothenburg University, Sweden, and incubated overnight at 37°C for CS3 and at room temperature for GM1 ganglioside / LTB, respectively. The following day, the plates were washed with PBS and blocked using a 1% BSA/PBS solution. After blocking, the plates were washed with 0.5% Tween-PBS. Samples diluted in BSA-PBS-Tween buffer were serially diluted across the plate and incubated for 90 minutes at room temperature. Following incubation, the plates were washed, and anti-human IgG-HRP (Jackson 309-035-006) was added and incubated for another 90 minutes. After a final wash, the substrate (TMB) was added, followed by the stop solution (1M sulphuric acid), and absorbance was measured at 490 nm using a BioTek ELISA plate reader and analysed using Gen 5 software.

Antibody titres were determined by plotting absorbance values (linear scale) against sample dilutions (log scale). The titre was defined as the interpolated dilution that produced an absorbance value of 0.4 above the background.

Samples collected during the three time points (baseline, day 22, and day 97) from each child were all analysed on the same plate to prevent the effect of variations in testing conditions. A reference sample was also included in each run to ensure consistency and repeatability across runs.

### Statistical Analysis

Differences in mean avidity indices between time points were assessed using paired t-tests with two-tailed *p* values, and statistical significance was set at *p* < 0.05. All statistical analyses were performed in R version 3.14.

Spaghetti plots were used to visualize individual trajectories of antibody avidity indices across visits, focusing on participants who demonstrated increases between any two time points.

Naïve status was assessed using both titre-based and avidity-based definitions. For titres, empirical cut-offs were set at the 20th percentile of baseline distributions for each antigen and isotype. Avidity-defined naivety was defined as an avidity index (AI) < 0.5, consistent with thresholds applied in studies of measles[15], malaria [16], and influenza [17].

Receiver operating characteristic (ROC) analysis was conducted to evaluate the ability of avidity indices to discriminate titre-defined naïve from non-naïve participants. The area under the ROC curve (AUC) was used as a measure of classification accuracy, and the optimal cut-off was determined using Youden’s J statistic. ROC analyses have been widely applied in infectious disease serology to define diagnostic or immunological thresholds [15, 18]. In this study, ROC analyses were performed separately for CS3 and LTB.

## Results

### Demographic Characteristics of participants

We assessed the avidity of anti-CS3 and anti-LTB IgG in plasma samples from 60/75 (80%) participants aged 6 to 23 months who were vaccinated with ¼ dose of the ETVAX vaccine with dmLT adjuvant that met the criteria (i.e. received a quarter dose of ETVAX^®^ and attended all study visits, ensuring blood collection at all three time points (Days 1, 22, and 97) were identified). Seventeen (17) of these were from cohort B and 43 were from cohort C (**Fig 1**).

**Table 1** shows selected demographic characteristics of the participants included in this analysis. Two-thirds were males (66.7%) and a third were females. The mean age was 16.1 months in cohort B, and 7 months in cohort C. The mean weight was 9.3kg for the older children (10-23 months) and 8 kg for the younger children (6-9 months). Nutritional status measured using the mid upper arm circumference (median = 14.5cm, (range: 12.5-18.0)) indicated that the cohort was healthy with no acute malnutrition.

**Table 1:** Participant demographics for cohorts B and C that received ¼ dose of ETVAX.

| Cohort | Cohort B (Children 10–23 months)<br>N=20 | Cohort C (Children 6–9 months)<br>N=55 | Total Participants<br>N=75 |
| --- | --- | --- | --- |
| **Gender** |  |  |  |
| Female | 7 (35.0%) | 18 (32.7%) | 25 (33.3%) |
| Male | 13 (65.0%) | 37 (67.3%) | 50 (66.7%) |
| **Age (months)** |  |  |  |
| Mean (SD) | 16.1 (3.43) | 7.0 (1.15) | 9.5 (5.09) |
| Median (Min, Max) | 16.0 (10, 23) | 7.0 (6, 9) | 8.5(6, 23) |
| **Height (cm)** |  |  |  |
| Mean (SD) | 74.8 (4.82) | 66.8 (2.98) | 69.1 (5.24) |
| Median (Min, Max) | 73.6 (64, 86) | 67.0 (59.2, 73.0) | 70.0 (59.2, 86.0) |
| **Weight (Kg)** |  |  |  |
| Mean (SD) | 9.3 (0.81) | 8.0 (1.11) | 8.4 (1.14) |
| Median (Min, Max) | 9.3 (7.8, 11.0) | 8.1 (5.7, 10.5) | 8.6 (5.7, 11.0) |
| **Mid Upper Arm Circumference (cm)** |  |  |  |
| Mean (SD) | 14.3 (0.63) | 14.6 (1.28) | 14.5 (1.10) |
| Median (Min, Max) | 14.3 (13.2, 15.5) | 14.5 (12.5, 18.0) | 14.4 (12.5, 18.0) |

### CS3 avidity

The baseline mean avidity index was 0.7 (SD=0.22; 95% CI: 0.64-0.75) for anti-CS3 IgG and remained unchanged (mean=0.7, SD=0.23) after the second dose and decreased (mean =0.6), SD=0.22; 95% CI:0.54-0.65) after the third dose (**Fig 2 and Table 2**). There was a statistically significant decline (p=0.00046) on comparison of the difference in means between baseline and visit 7 (7 days after the third dose) and between visit 5 and visit 7 (p=0.00022).

**Fig 2:**
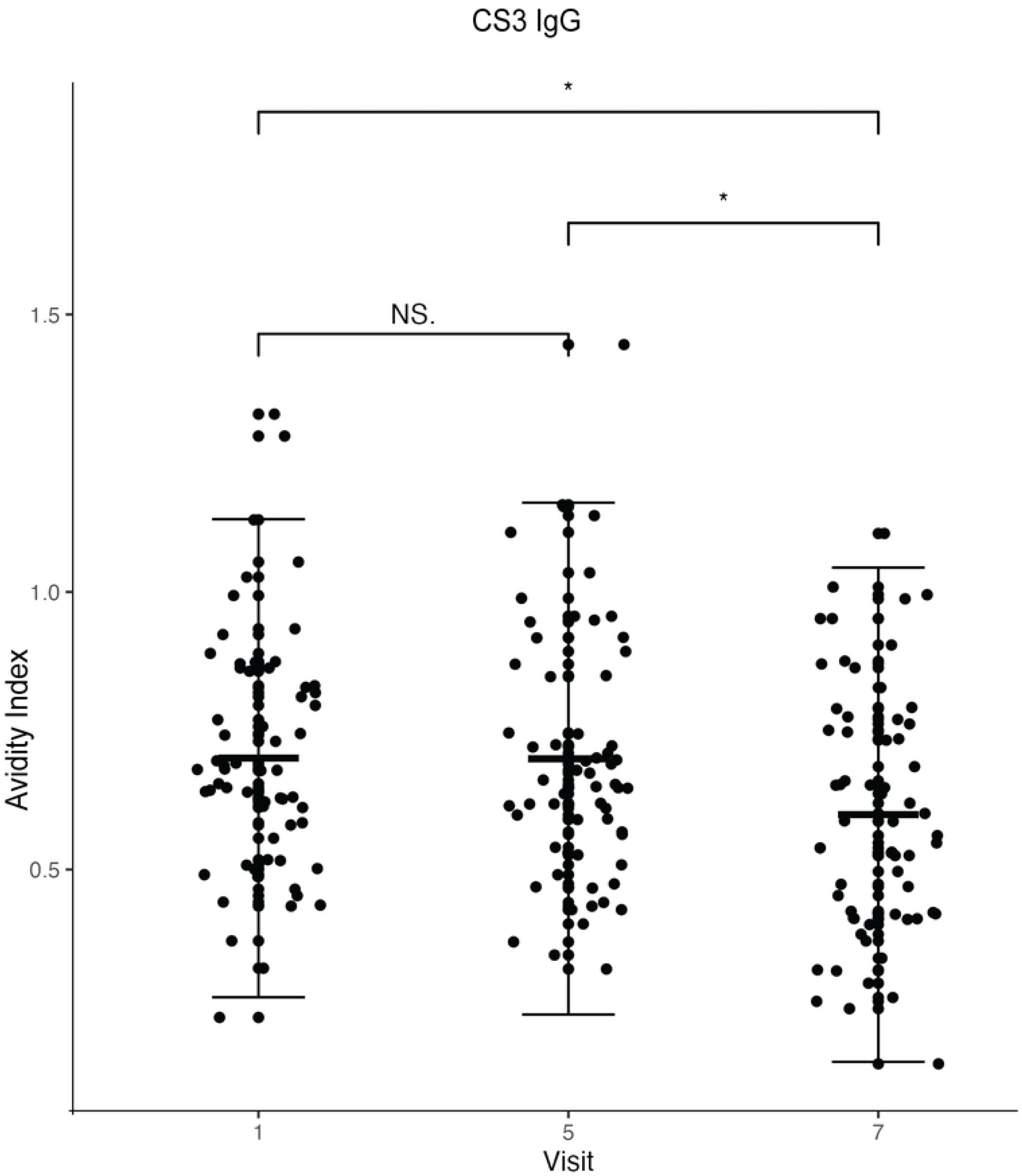
Plot of plasma anti-CS3 IgG antibody avidity index against the visit (visit 1= baseline, visit 5=7 days after the second dose and visit 7= 7 days after the third dose). The dots represent each participant (n=60). *The Wilcoxon matched pairs signed rank test was used for statistical comparisons; *P < 0.05. The central horizontal line shows the mean while the vertical line error bars show the standard deviations*.

**Table 2:**
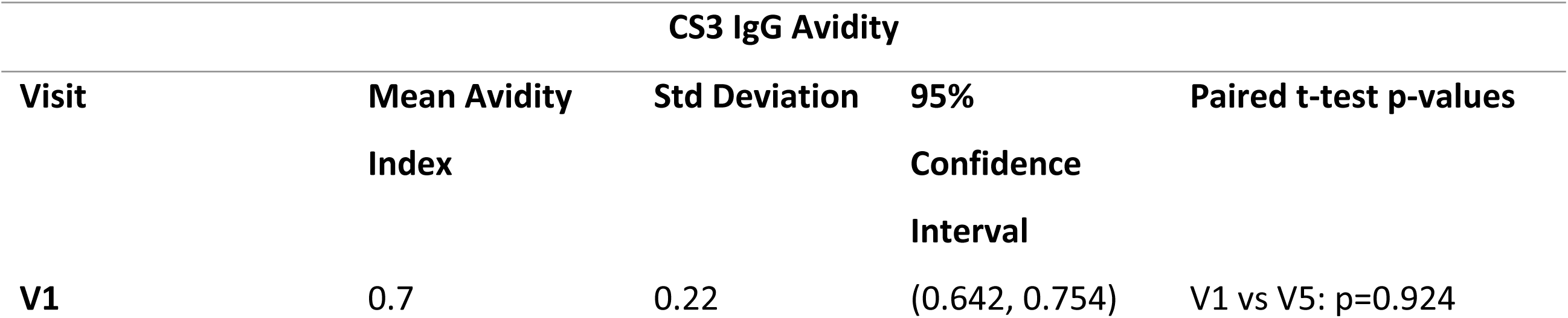
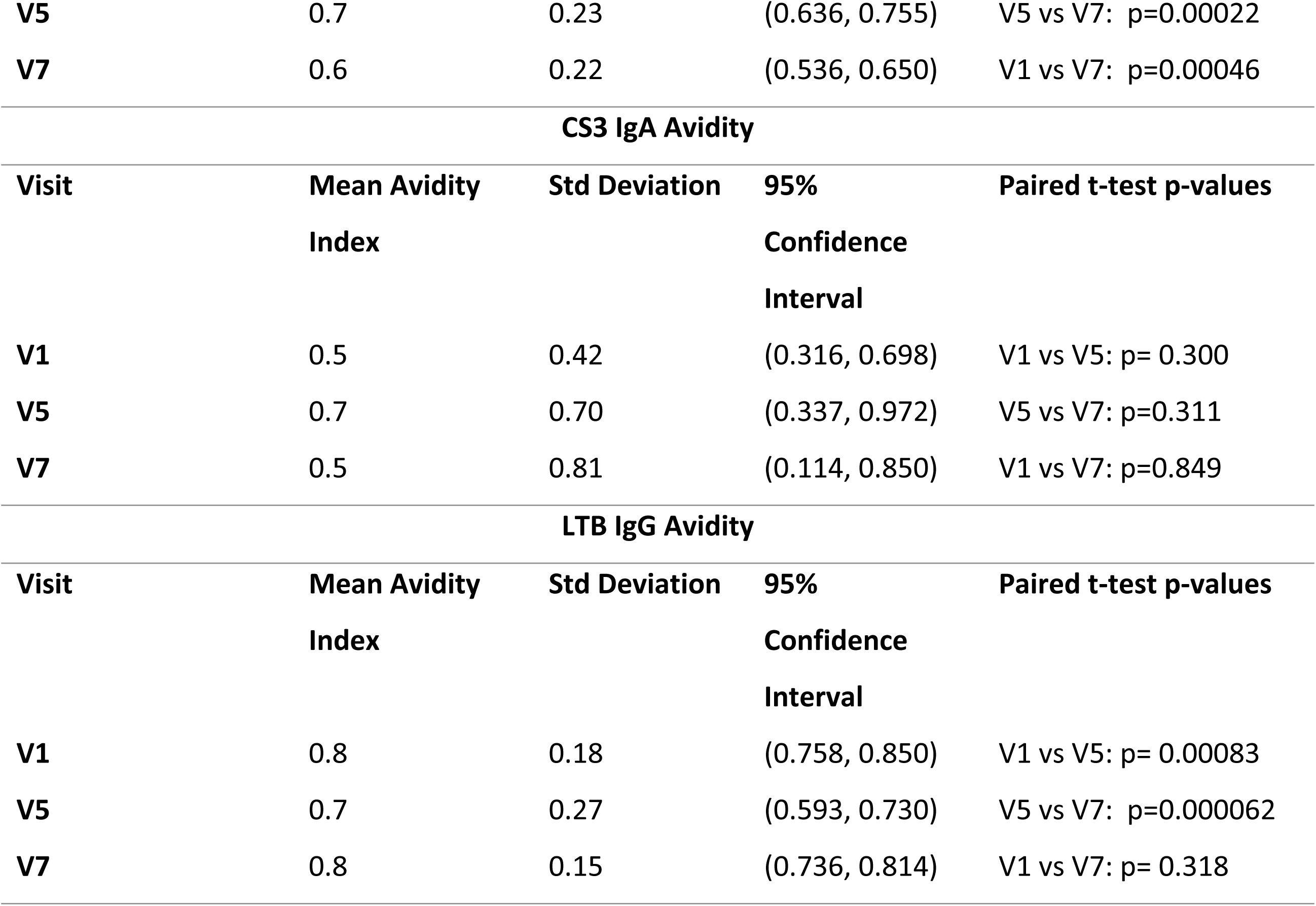
Mean AI values at the three timepoints.

We also analysed a few randomly selected samples (n=20) for plasma anti-CS3 IgA avidity, due to limited reagents (**Fig 3**). The pattern in the changes in the mean values at the different time points was different from that observed for IgG. There was an increase from 0.51 to 0.65 (V1 to V5 respectively) and a drop to 0.48 at V7 (**Table 2**). These changes were however not statistically significant (p=0.300, 0.311 and 0.849 respectively).

**Fig 3:**
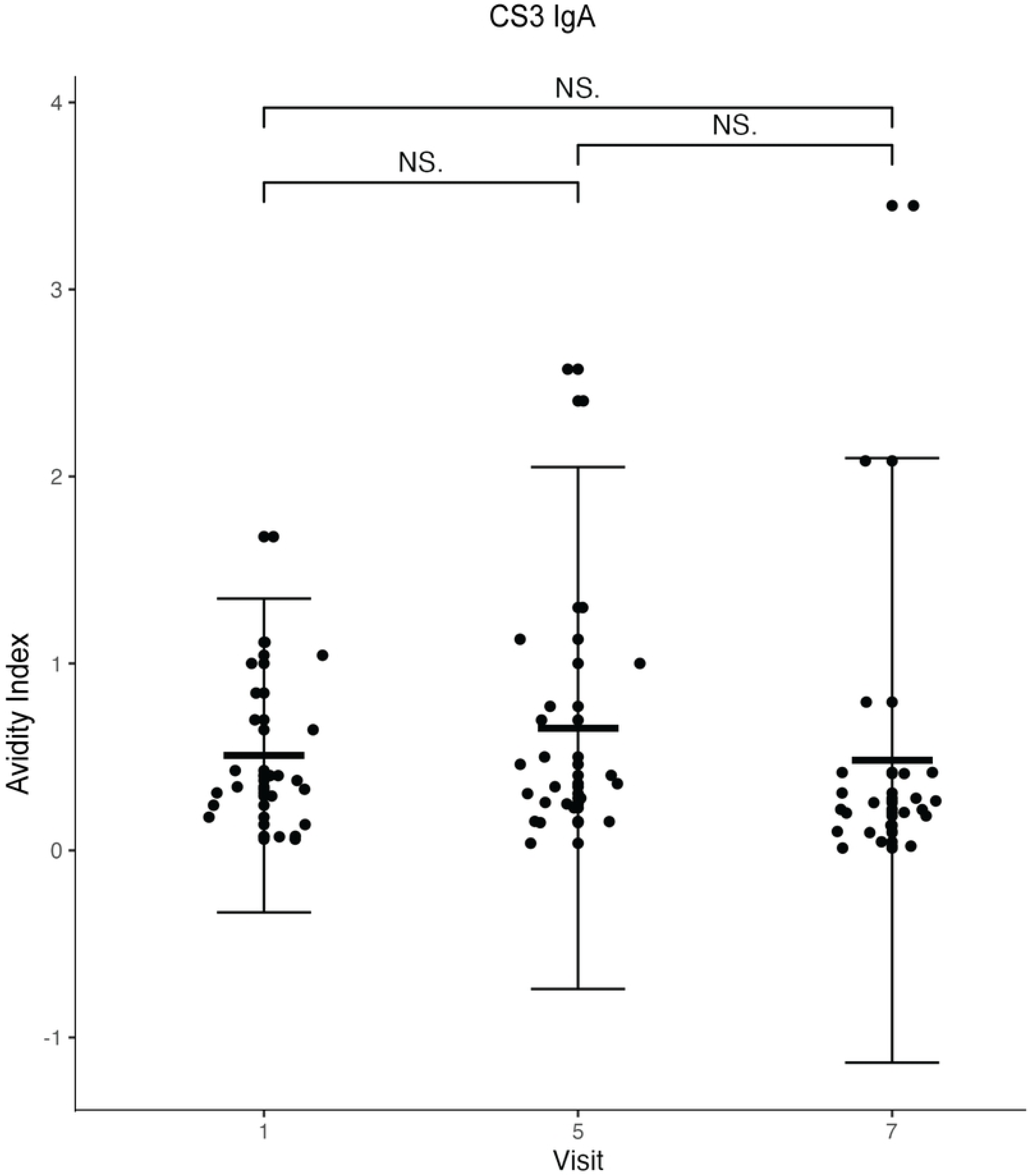
Plot of plasma anti-CS3 IgA antibody avidity index against the visit (visit 1= baseline, visit 5=7 days after the second dose and visit 7= 7 days after the third dose). The dots represent each participant (n=20). *The Wilcoxon matched pairs signed rank test was used for statistical comparisons; *P < 0.05. The central horizontal line shows the mean while the vertical line error bars show the standard deviation*.

### LTB avidity

There was a statistically significant decrease in the IgG antibody avidity following the second dose (p=0.00083), and a significant increase from the second to the third dose (p=0.000062) (**Fig 4 and Table 2**). However, the increase after the third dose was not significantly higher than baseline (p=318).

**Fig 4:**
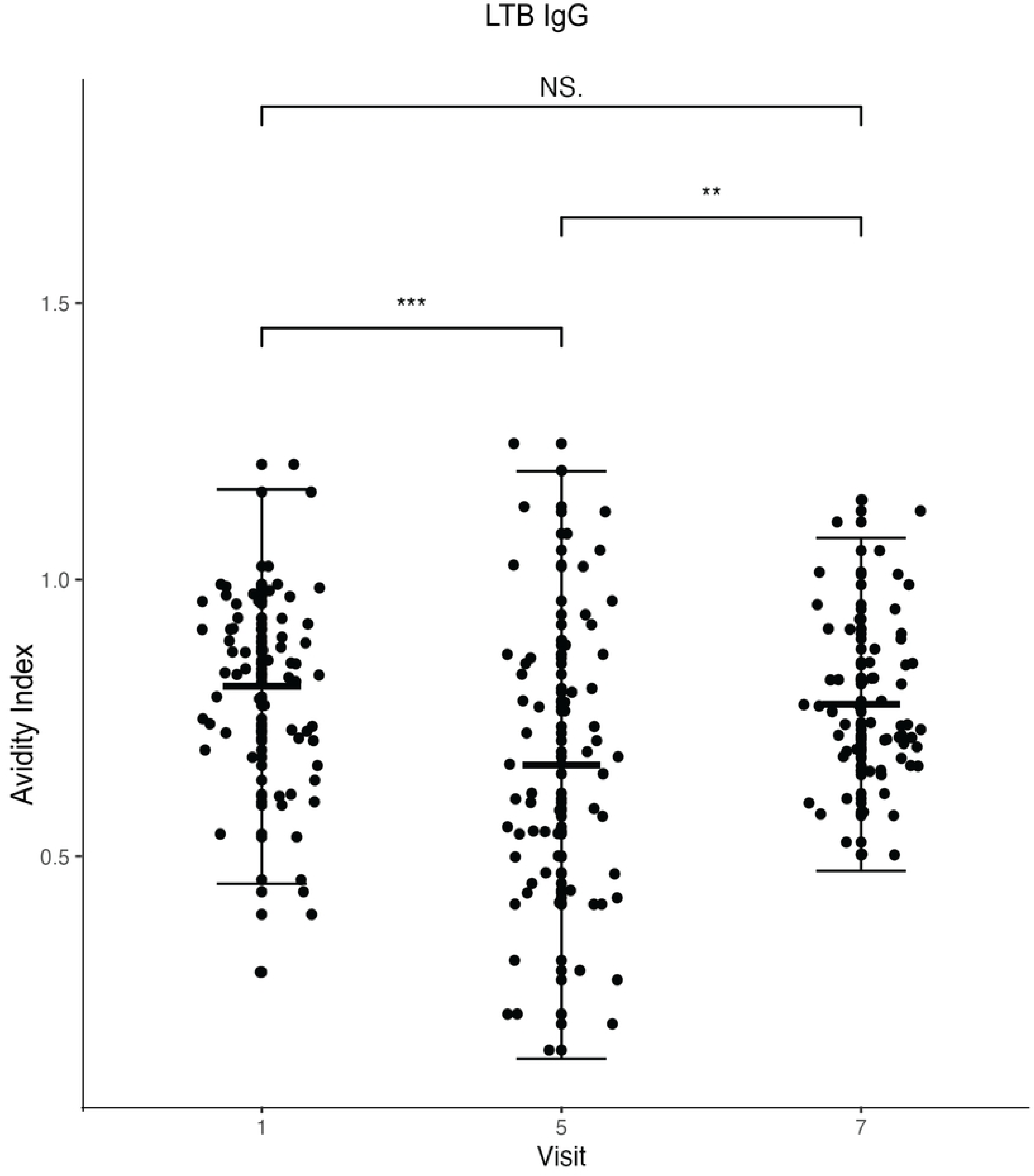
Plot of plasma anti-LTB IgG antibody avidity index against the visit (visit 1= baseline, visit 5=7 days after the second dose and visit 7= 7 days after the third dose). The dots represent each participant (n=60). *The Wilcoxon matched pairs signed rank test was used for statistical comparisons; *P < 0.05. The central horizontal line shows the mean while the vertical line error bars show the standard deviation*.

### Participants with increases in avidity

To test our hypothesis that avidity increases with increasing vaccine doses, samples were stratified using two criteria; (A) those with a net avidity increase in avidity from Visit 1 (V1) to Visit 7 (V7), and (B) those strictly increasing avidity at Visits 1, 5, and 7 (**Fig 5**). Panel A shows participants with net increase in avidity index (that is, the AI at V7 is greater than that at V1). In this group, we observed inter-individual variability, particularly in the response in the CS3- specific IgA where the subset had extensive avidity index increments by V7. In contrast, CS3- and LTB-specific IgG responses demonstrated more modest net increases, with several individuals reaching peak AI values at the intermediate timepoint (V5), suggestive of heterogeneous kinetics of affinity maturation.

**Fig 5:**
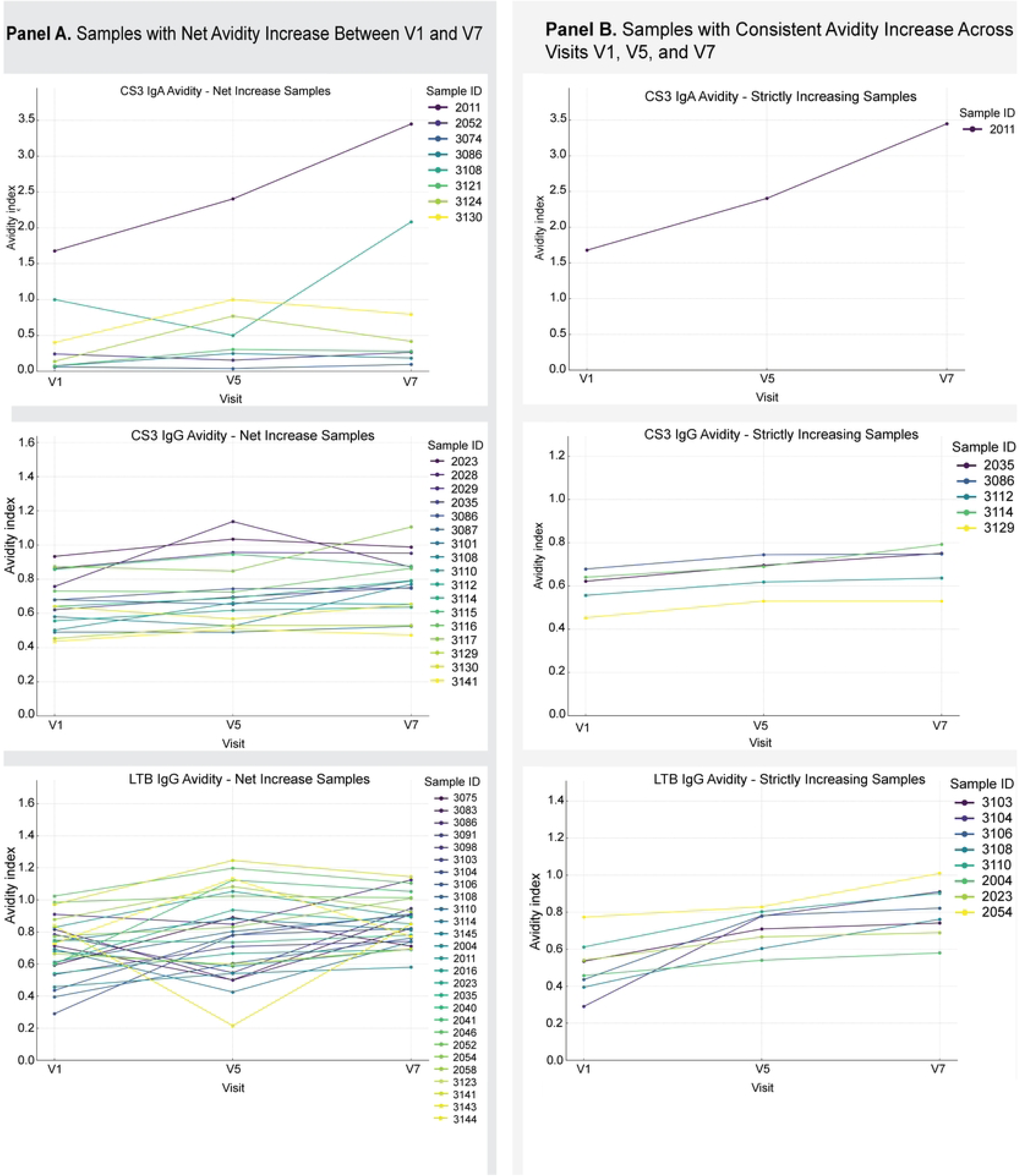
Longitudinal avidity index trajectories for samples with net or consistent avidity increases across visits V1, V5, and V7. **Panel A** shows samples with a net increase in avidity index between visits V1 and V7 across three antibody datasets: CS3 IgA (top), CS3 IgG (middle), and LTB IgG (bottom). Each line represents an individual sample’s avidity trajectory over time. Colors correspond to individual sample IDs as indicated in the legends. **Panel B** shows samples with strictly increasing avidity indices across all three visits (V1, V5, and V7).

Panel B focused on participants whose antibody avidity indices increased consistently across all visits. The number of participants fulfilling this condition was very small and included one participant (sample ID 2033) whose level of CS3-specific IgA avidity had particularly significantly increased. For CS3, 5/60 and for LTB, 8/60 participants met these criteria. Although the magnitude of these increases in avidity was generally small, some participants showed a steady rise at every timepoint. This steady pattern was uncommon. While it’s normal to see overall improvement, it’s rare to see a smooth, step-by-step increase. When this happens, it might be due to unique immune system traits or previous exposures to similar germs

### Correlation of avidity and antibody titre data

We performed correlation analyses for avidity and titres to see the relationship between antibody titres and antibody avidity (**Fig 6**). For CS3 IgA, we observed generally weak to moderate relationships between antibody titres and avidity indices across visits V1, V5, and V7. None of the correlations reached statistical significance at the 0.05 level. The scatter plots reveal some variation and no clear linear trend, suggesting that avidity maturation in IgA antibodies may not strongly parallel titre changes at these time points. Analyses for CS3 IgG revealed a significant correlation was observed at visit V7 with a moderate negative Pearson correlation (r = -0.37, p = 0.004), indicating that at this late time point, higher titres corresponded with somewhat lower avidity indices or vice versa. Visits V1 and V5 did not show significant correlations. LTB IgG showed moderate significant negative correlation (Pearson r = - 0.42, p = 0.00074) at V1, similar to CS3 IgG V7, implying early inverse relationships between titre and avidity. No significant correlations were found at visits V5 and V7

**Fig 6:**
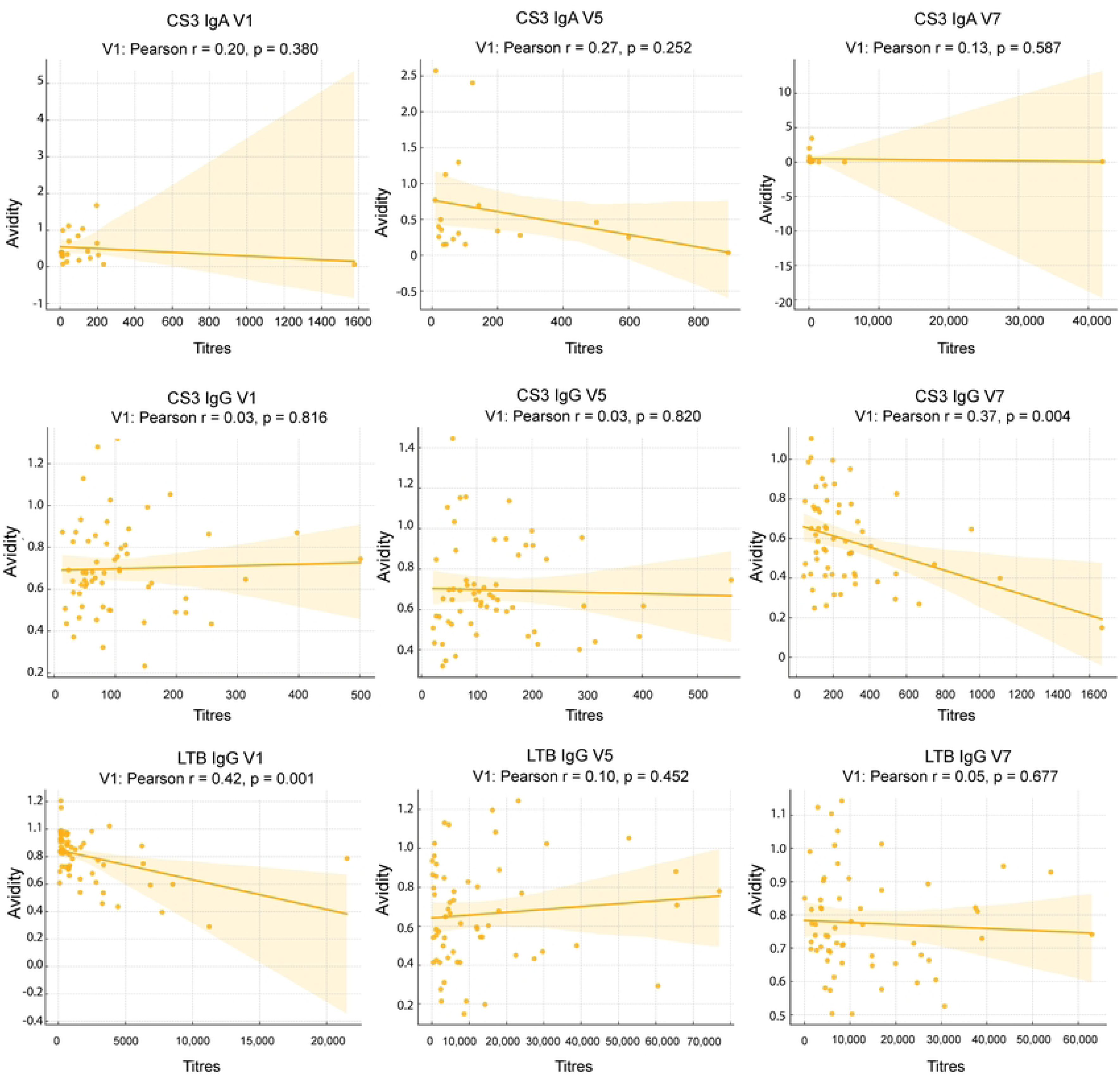
Correlation between antibody titres and avidity indices across visits for CS3 IgA, CS3 IgG, and LTB IgG. Scatter plots show the relationship between antibody titres (x-axis) and avidity indices (y-axis) for visits V1, V5, and V7. Each panel includes a linear regression line with 95% confidence interval shaded in orange. Pearson correlation coefficients (r) and associated p-values are indicated above each plot. Significant correlations highlight temporal dynamics in antibody quality and quantity

### Naïve classification

Baseline naïve status of participants was assessed using both titre-based and avidity-based definitions as shown by quadrant plots in **Fig 7**. Empirical titre cut-offs were derived from the 20th percentile of baseline titres (CS3 IgG ≈44, CS3 IgA ≈16, LTB IgG ≈203, LTB IgA ≈35), while avidity-defined naivety was classified as AI <0.5.

**Fig 7.**
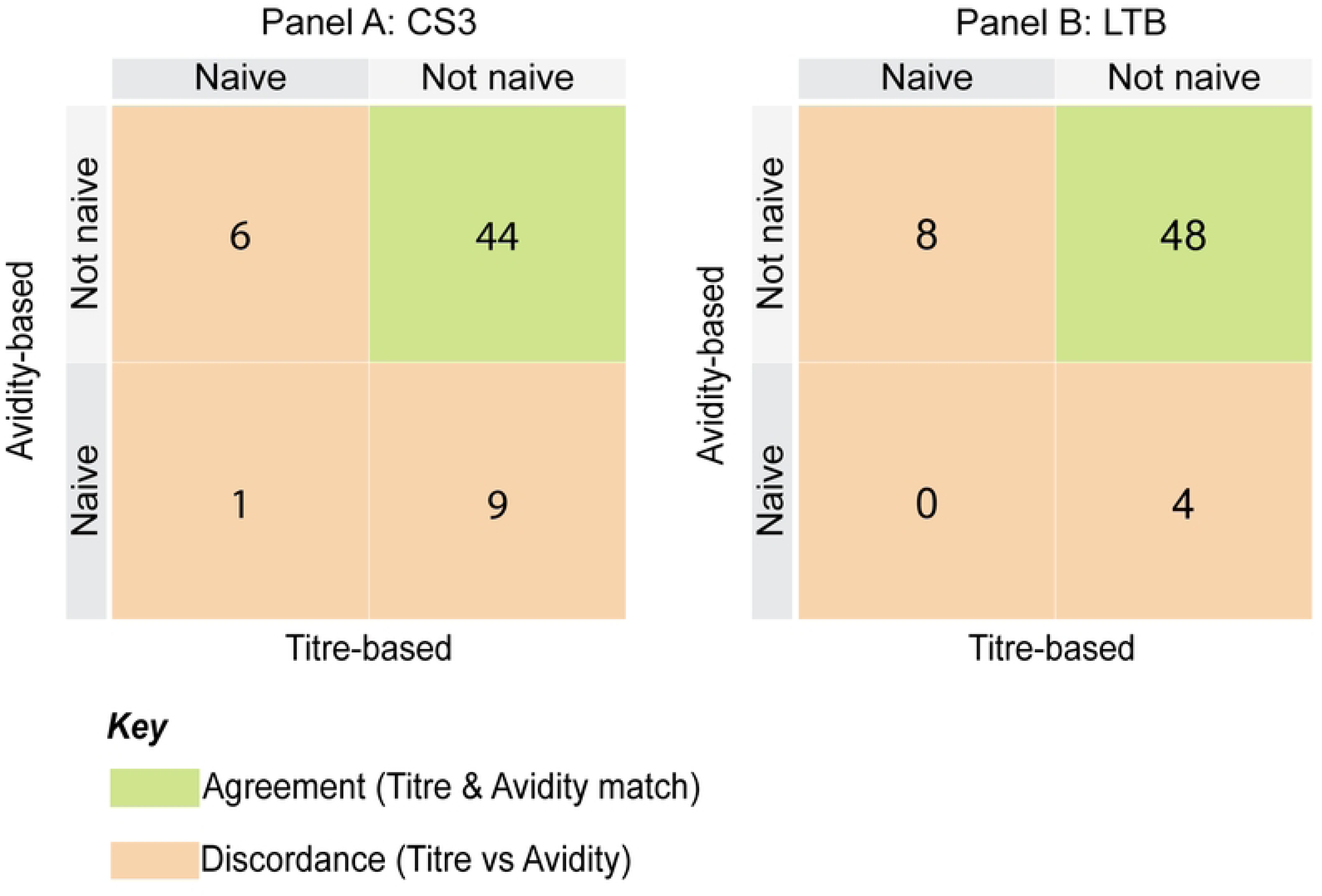
Comparison of titre- and avidity-based naïve classification for CS3 and LTB. Quadrant plots display naïve status classification determined using titre-based cut-offs (20th percentile of baseline titres) and avidity-based cut-offs (AI < 0.5) outcomes for CS3 (Panel A) and LTB (Panel B). Blue quadrants indicate agreement between methods, while orange quadrants highlight discordant classifications.

For CS3, 44 participants were consistently non-naïve by both methods. Nine participants (15%) had titres but low avidity, indicating functional naivety, while six (10%) were naïve by titres but non-naïve by avidity, consistent with waned titres and preserved antibody quality. Only one child (2%) was naïve by both criteria.

For LTB, 48 participants were non-naïve by both methods. Four (7%) had titres but low avidity, while eight (13%) were naïve by titres but non-naïve by avidity. No participants were naïve by both definitions.

### ROC analysis

Receiver operating characteristic (ROC) analysis was performed to evaluate the ability of avidity indices to discriminate titre-defined naïve from non-naïve participants (**Fig 8**).

**Fig 8.**
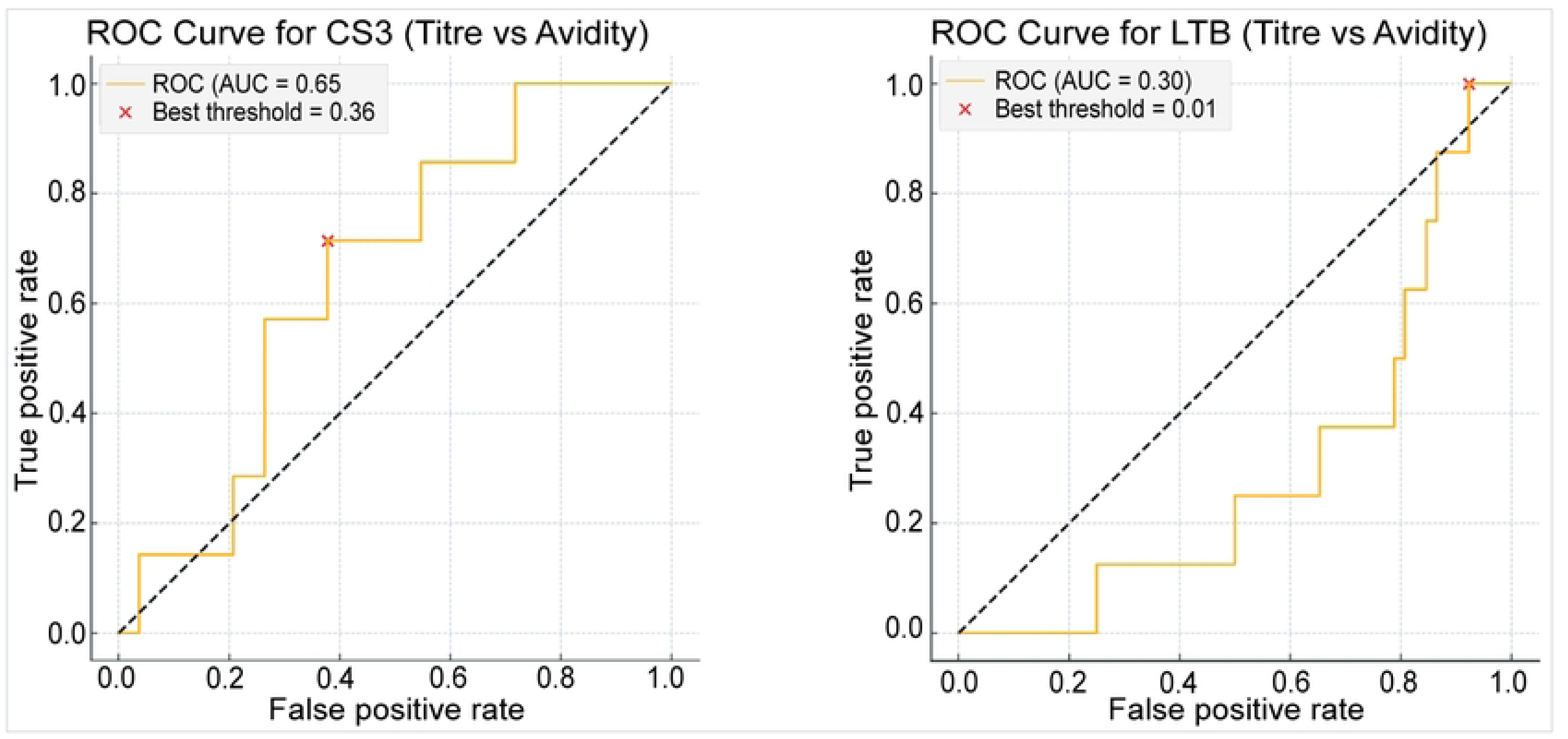
Receiver operating characteristic (ROC) analysis of antibody avidity for CS3 and LTB. Panel A shows the ROC curve for anti-CS3 IgG avidity (AUC = 0.65), with the optimal cut-off identified at an avidity index of 0.36 (red cross). Panel B shows the ROC curve for anti-LTB IgG avidity (AUC = 0.30), where the optimal cut-off was 0.01. The diagonal dashed line represents random classification.

For CS3, the ROC curve showed moderate discrimination (AUC = 0.65), with the optimal avidity cut-off identified at 0.36, slightly below the conventional 0.5 threshold. In contrast, LTB avidity demonstrated poor discriminatory ability (AUC = 0.30), with an optimal cut-off of 0.01, consistent with the near absence of naïve participants for this antigen.

## Discussion

We assessed the antibody avidity of plasma IgG in children aged 6-23 months participating in a dose selection phase 1b trial assessing the safety and tolerability of the ETVAX vaccine in Zambia. This study builds upon our previous work in which we presented data on the IgA titres to CFA/I, CS3, CS5, CS6 and LTB and anti-LTB IgG titres [13]. We reported significant increases in antibody titres following vaccination which increased with increasing vaccine doses [13].

Our findings demonstrated that participants had detectable avidity to the ETEC antigens LTB and CS3 prior to vaccination. The detectable baseline avidity mirrored baseline antibody titre findings previously described for this cohort which may be attributed to pre-existing immunity from natural ETEC exposure [13]. Our previous work, along with data from other studies, shows that in ETEC-endemic areas like Zambia, diarrhoea caused by ETEC is common in children under five years. A study from Peru also found that infections can start soon after birth. [5, 19–21].

We observed minor changes in antibody avidity index after vaccination with differing response patterns for the two antigens (CS3 and LTB) assessed. For anti-CS3 IgG, we observed a statistically significant overall decrease in AI after the third dose compared to baseline. For anti-LTB IgG, a transient drop in AI was observed after the second dose, followed by an increase after the third dose, which was not significantly higher than baseline. This is consistent with the findings from studies of other pathogens such as malaria, in which they observed that some antigens tend to induce affinity maturation better than others and affinity maturation doesn’t occur at the same rate and doesn’t reach the same level for all antigens [12]. For ETEC, studies show that more robust immunity tends to develop against homologous antigens (homotypic immunity) compared to heterotypic immunity, therefore, the antibody avidity patterns observed in this study may be attributed to the influence of prior antigen exposure[8, 22].

Our findings challenge the expectation that repeated vaccination results in increasing avidity, an understanding rooted in the classical understanding that repeated antigen exposure results in high antibody affinities to pathogens and consequently higher antibody avidity[16]. Several possible explanations may account for this. The endemic nature of ETEC in Zambia results in repeated natural exposure to the pathogen, providing continuous antigenic stimulation, and thereby maintaining high avidity antibodies even before vaccination [8, 23].

Repeated vaccination may lead to a ceiling effect, where additional doses do not significantly enhance antibody avidity. This phenomenon, known as “antigen trapping”, occurs when pre-existing antibodies and memory B cells from prior infections or vaccinations rapidly bind to the new antigen. As a result, less antigen remains available to stimulate naïve B cells, limiting further avidity maturation [24]. This effect has been observed in studies where previously infected individuals exhibited high baseline avidity but showed minimal increases with subsequent vaccine doses. In contrast, naïve individuals demonstrated greater avidity maturation over time [25].

Secondly, the quarter dose of ETVAX^®^ used in this study, although found to be immunogenic and inducing very good antibody titres might not be optimal for inducing significant changes in antibody avidity in children in endemic areas like Zambia [13] . Therefore, future studies could explore different dosing regimens or formulations to better understand their impact on avidity maturation and a killing assay.

In another study, Yam et al. reported unusual patterns of IgG avidity in young children following two doses of an adjuvanted H1N1 flu virus vaccine [17]. Some primed children showed a constant AI from day 0 to day 42, suggestive of a fully mature IgG avidity at the time of vaccination, compared to naive children whose AI increased after the first dose but declined steeply after the second dose, indicative of a more dynamic avidity response[17].

We observed that a small subset of participants exhibited measurable increases in antibody avidity over time, both in terms of overall net gain and consistent, strictly increasing trends (Figure 5). These individual variations may reflect differences in immunological priming, timing of prior infections, or host-specific factors affecting B-cell maturation. Yam et al also observed this, that patterns of the antibody avidity were different for each individual child in their study and highlighted the need to report individual avidity patterns apart from aggregate data only as this provides more insight into the immune response of each individual participant, considering their individual immune dynamics [17].

Correlation analyses between antibody titres and avidity showed that the humoral immune response can be quite complex. The negative correlations observed at some visits indicates that high antibody titres are not always associated with greater avidity. Similar observations have been reported in studies of malaria. High affinity antibodies are a product of affinity maturation which occurs when low-affinity B cells undergo immunoglobulin (Ig) somatic hypermutation (SHM), clonal expansion, and affinity-based selection within the germinal centre (GC) following exposure to antigens through infection or vaccination[10]. Ssewanyana et al. have suggested that in settings with constant antigen exposure, such as malaria endemic areas, affinity maturation may be impaired, potentially due to two non-mutually exclusive mechanisms. First, constant exposure to an antigen may hinder the maturation or persistence of high-avidity antibodies through various mechanisms. Second, a recent infection may introduce low-avidity antibodies into circulation, thereby reducing the overall proportion of high-avidity antibodies [16]. They observed an inverse relationship between the avidity of anti-malaria antibodies and transmission intensity among children under five and found that children in high-transmission areas had low-avidity antibodies despite having high antibody levels [16]. Additionally, they noted that avidity to two *Plasmodium falciparum* antigens was significantly lower in these high-transmission areas [16].

To further assess baseline immune maturity, we classified children using both titre-based and avidity-based definitions of naïve status. Empirical titre cut-offs (20th percentile of baseline titres) were combined with a conventional avidity threshold (AI <0.5). For CS3, most children were classified as non-naïve by both measures, but one child was naïve by both, nine (15%) had titres but low avidity (“functional naivety”), and six had low titres but retained high avidity (“waned titres with memory”). For LTB, no child was naïve by both measures, and 12 discordant cases were observed. Similar frameworks have been applied in measles [15], mumps [26], and malaria [18], where avidity was used to reveal functional naivety despite measurable titres.

The ROC analysis further evaluated this framework by testing the discriminatory performance of avidity indices against titre-based definitions. For CS3, the AUC of 0.65 suggests modest ability of avidity to refine naïve classification, consistent with findings from measles [15] and malaria vaccine studies [18]. The identification of an optimal cut-off (0.36), slightly below the conventional 0.5 threshold, underscores the need for context-specific thresholds in endemic settings. In contrast, the poor performance of LTB avidity (AUC = 0.30) reflects the near-universal prior exposure to LT antigens in young children, limiting the ability of either titres or avidity to separate naïve from non-naïve individuals. This pattern mirrors observations from endemic viral infections where ROC analysis failed to provide meaningful separation [17].

Together, these results demonstrate both the utility and limitations of ROC-derived thresholds in refining immunological classification in endemic populations.

Understanding antibody avidity is crucial for vaccine design and dose optimization, as high-avidity antibodies are more effective at neutralizing pathogens and providing long-term immunity [11, 27]. Our results highlight the importance of evaluating both titres and avidity to fully characterise the immune responses to vaccines and natural infections. The minimal changes in avidity observed in this study show the need for further investigation, particularly into the role of T-helper cells and other immune components in supporting avidity maturation. Correlating antibody avidity with T-cell responses could provide a more comprehensive understanding of the immune mechanisms involved and inform strategies to enhance vaccine efficacy.

To our knowledge, this is the first study to assess antibody avidity to the ETVAX vaccine in children within our setting. However, the interpretation of our findings is limited by the lack of comparable studies. Additionally, the small sample size restricted our ability to analyse factors influencing antibody development. The statistical significance and strength of correlations also depend heavily on sample size. Another limitation was the insufficient availability of reagents and resources, which prevented a comprehensive assessment of the anti-IgA avidity. The sampling schedule may be another aspect to consider in that avidity maturation may occur beyond our final sampling point (day 97), and our study may have missed later affinity increases.

Future research should include longitudinal assessing antibody avidity alongside titres and T cell responses to gain insights into the durability of vaccine-induced immunity. Our naïve classification and ROC data suggest that special attention should be given to baseline immune maturity, since titres alone may overestimate exposure history in endemic settings. Expanding the study cohort to include individuals with no prior exposure ETEC, such as neonates or participants from non-endemic regions, could help differentiate the effects of natural immunity from vaccine-induced responses. Additionally, incorporating functional antibody assessments, such as neutralization assays, would complement avidity measurements and offer a more complete evaluation of vaccine efficacy.

## Conclusion

This study highlights the complexity of antibody responses to vaccination in children from ETEC- endemic settings. While ETVAX^®^ induced strong antibody titres, antibody avidity showed limited changes after vaccination, with notable inter-individual variability. Naïve classification analyses demonstrated that titres alone may overestimate prior immune maturity, and ROC analysis indicated that avidity provides additional discriminatory value, though with context-specific limitations. Together, these findings underscore the importance of assessing both antibody quantity and quality when evaluating vaccines in endemic populations.

## Acknowledgements

We thank the children and families who participated in the ETVAX trial whose samples were used in this study. Grateful to Mr. Kelvin Mwangilwa from ZNPHI for statistical insights. And to the members of the Basic Science and Immunology department at CIDRZ for the moral support.

